# Administration of short-chain fatty acids reduces bacterial translocation to peripheral organs and the brain, preserves gut barrier integrity, and mitigates brain injury following stroke

**DOI:** 10.1101/2025.11.03.686233

**Authors:** Julia Castillo-González, Seyedeh Maryam Mousavi, José Luis Ruiz, Roméo Carre, Valentine Bressoud, Lara Buscemi, Melanie Price, Solenne Vigne, Caroline Pot, Lorenz Hirt

**Affiliations:** Service of Neurology, Department of Clinical Neurosciences, Lausanne University Hospital and University of Lausanne, Lausanne, Switzerland; Department of Fundamental Neurosciences, University of Lausanne, Lausanne, Switzerland; Functional Genomics Center Zurich, ETH Zurich and University of Zurich, Zurich, Switzerland; Swiss Institute of Bioinformatics, Amphipôle, Quartier UNIL-Sorge, Lausanne, Switzerland; Laboratories of Neuroimmunology, Service of Neurology, Department of Clinical Neurosciences, Lausanne University Hospital and University of Lausanne, Lausanne, Switzerland

**Keywords:** stroke, bacterial translocation, post-stroke infections, short-chain fatty acids, gut metabolites, brain-immune-gut axis, gut barrier

## Abstract

**Background:** Stroke remains a leading cause of death globally, with post-stroke infections (PSIs) representing its main life-threatening complication. While traditionally attributed to nosocomial infections or medical procedures, emerging research has revealed that PSIs primarily originate from bacterial translocation following gut barrier disruption. Yet, the phenomenon of bacterial translocation remains poorly characterized. Critical knowledge gaps persist regarding the exact spatiotemporal dynamics of this event, and whether bacterial dissemination actively drives stroke pathogenesis and worsens neurological outcomes. Consequently, the development of targeted therapies remains severely limited. In this context, short-chain fatty acids (SCFAs) have emerged as key regulators of the brain-gut-immune axis, exerting neuroprotective and anti-inflammatory properties. However, their potential therapeutic modulation of bacterial translocation has remained unexplored.

**Methods and Results:** Using a middle cerebral artery occlusion model in mice, we confirmed that stroke induces systemic bacterial dissemination to the liver, lungs, and spleen, and provided novel evidence of the presence of bacteria in the heart, brainstem, and ischemic lesion core. Interestingly, systemic bacterial load correlated with worse neurological outcomes but not with ischemic volume. Furthermore, we described for the first time the spatiotemporal kinetics of this process, demonstrating that translocation initiated as early as 6 hours post-stroke, peaked between 24 hours and 4 days, and had almost completely resolved systemically by 7 days, while strikingly persisting in the central nervous system. Importantly, SCFA treatment reduced both systemic and cerebral bacterial translocation, preserved gut barrier integrity, and reversed post-stroke immunosuppression. SCFAs also reduced infarct volume and blood-brain barrier leakage, ultimately improving survival rates.

**Conclusion:** These findings provide crucial insights into the complex process of bacterial translocation and highlight SCFAs as a promising therapeutic strategy for both the ischemic event and its most lethal, yet least understood, complications.

## Introduction

Ischemic stroke results from the occlusion of a brain artery, which provokes irreversible tissue injury and long-term sequelae. Neuroinflammation, blood-brain barrier (BBB) breakdown, and immune dysregulation are major hallmarks of stroke pathogenesis^1^. Recent studies have highlighted the pivotal role of the brain-immune-gut axis (*i.e.,* complex multidirectional communication between the nervous system, the immune system, and the gut). Stroke dysregulates this axis, leading to gut dysbiosis (*i.e.,* alterations in microbiota composition) and increased gut barrier permeability, which correlate with impaired recovery and a worse stroke outcome^2^. Despite improvements in recanalization treatments, stroke is the second leading cause of death worldwide and the main cause of disability in adults^3^. In this context, post-stroke infections (PSIs) are the main life-threatening complications (affecting 30-45% of all patients and accounting for 20% mortality), with bacterial pneumonia being the most frequent within the first week after stroke^4^. PSIs have traditionally been attributed to medical procedures or nosocomial infections^5^. However, a groundbreaking study reported that over 70% of the bacteria detected in patients suffering from PSIs were common commensal bacteria from the intestinal tract^6^. This suggests that PSIs largely result from bacterial translocation of host gut strains, disseminating after gut barrier disruption. Notably, the profound and complex immunosuppression that follows the initial pro-inflammatory response after stroke increases susceptibility to these infections^1^. Despite this groundbreaking discovery, the causes, mechanisms, routes, spatiotemporal dynamics, and consequences of bacterial translocation remain poorly understood. Furthermore, while antibiotic prophylaxis has been investigated to prevent PSIs, three randomized controlled trials demonstrated no significant impact on stroke mortality or functional outcomes^7^. Overall, the lack of success of current therapies, likely due to the dysregulated immune response, increasing pathogen resistance to broad-spectrum antibiotics, and poor knowledge about bacterial translocation, underscores the need for a better understanding of the pathophysiology of PSIs.

Microbiota-derived compounds, commonly known as gut metabolites, are crucial regulators of the brain-immune-gut axis. Among these, short-chain fatty acids (SCFAs), small monocarboxylic acids resulting from the bacterial fermentation of polysaccharides, are of vital importance. The most abundant of these are acetate, propionate, and butyrate, typically found in a 60:20:20 ratio^8–10^. SCFAs play an important role in gut homeostasis and immune modulation^11,12^. Beyond these peripheral functions, SCFAs are also essential for brain homeostasis, microglial development, and neurotransmitter production^13,14^. Recent studies have identified low SCFA levels in both faecal content and blood of patients and experimental models of Parkinson’s disease, Alzheimer’s disease, and stroke, correlating with poorer prognoses^15–17^. Moreover, SCFAs have been reported to exert neuroprotective and anti-inflammatory effects in these disorders^9,13,18–20^. Although often overlooked in stroke, recent evidence indicates that SCFA administration or faecal transplant with SCFA-producing strains promotes functional recovery in mouse models of stroke by regulating the immune response^21–23^. Furthermore, beyond their neuroimmune properties, SCFAs have also shown antimicrobial effects in studies of colitis or wound infection, modulating pathogen growth and the immune response (reviewed in ^24^). However, the therapeutic potential of SCFAs in PSIs has not been previously elucidated, representing a significant knowledge gap. Therefore, in this study, we aimed to characterize the process of bacterial dissemination after stroke and evaluate its spatiotemporal dynamics in the MCAO model. Additionally, we sought to determine the therapeutic potential of SCFAs in mitigating this translocation, preserving gut barrier integrity, regulating the immune response, and ameliorating ischemic brain injury.

## Materials and methods

### Animals

All experimental procedures were conducted in strict accordance with the Swiss Federal Act on Animal Protection and the European Directive 2010/63/EU. The study protocol was reviewed and approved by the Vaud Cantonal Veterinary Office (Authorization VD2017.7d). Twelve-week-old male C57BL/6J mice (Charles River Laboratories, France) were used in all experiments. Animals were housed under controlled temperature and humidity conditions on a 12-hour light/dark cycle, with *ad libitum* access to food and water. All procedures were performed in compliance with the ARRIVE 2.0 guidelines for the design, analysis, and reporting of animal research. Whenever possible, the same animal was used for multiple tissue collections and experimental assessments, in accordance with the 3Rs principles to minimize animal use.

### Middle cerebral artery occlusion model

Cerebral ischemia was induced using the transient middle cerebral artery occlusion (MCAO) model^25–27^. Briefly, mice were anesthetized with isoflurane (1.5–2 % in 70 % N_2_O/30 % O_2_) using a facemask. Mice were administered buprenorphine subcutaneously at the beginning of the surgery and for post-surgical analgesia. Rectal temperature was monitored throughout surgery with a rectal probe and maintained at 37.0 ± 0.5 °C using a heating pad (FHC Inc.). Regional cerebral blood flow (rCBF) was monitored using laser Doppler flowmetry (Perimed AB, Sweden) by fixing a probe on the skull over the core area supplied by the middle cerebral artery (approximately 1 mm posterior and 6 mm lateral from bregma). Transient ischemia was induced by inserting a silicone-coated filament (701756PK5Re, Doccol Corp, Sharon, MA, USA) through the left common carotid artery into the internal carotid artery and advanced until it occluded the middle cerebral artery. The filament was left in place for 25 min before being withdrawn to allow reperfusion. Mice were excluded from the study if the rCBF did not drop below 20 % of the baseline during occlusion and/or did not rise above 50 % of the baseline within 10 min after filament withdrawal, as previously described^25–27^. The *sham* surgery procedure was identical, except that the filament was inserted through the common carotid artery into the internal carotid artery but not advanced into the arterial circle, ensuring no cerebral ischemia occurred (Suppl. Fig. 1A). This control was used to rule out bacterial presence resulting from surgical manipulation or filament insertion. After surgery, mice were housed with *ad libitum* access to food and water, with each cage containing water-softened chow.

To investigate the potential therapeutic role of SCFAs, mice received an intraperitoneal injection of either PBS (MCAO+PBS) or a mixture of sodium acetate (Sigma #S5636), sodium propionate (Sigma #P5436), and sodium butyrate (Sigma #303410) (MCAO+SCFAs) in a 6 mmol/kg body weight in a 60-20-20 ratio (*i.e.,* physiological human and mammalian molar ratios^28^) every 24 h from day 1 to day 3 post-stroke. Treatment assignments were performed randomly and in a blinded manner. Subsequent sample analyses were also conducted in a blinded way.

### Neuroscore

Mice were evaluated every 24 h using an adapted neuroscore scale ranging from 0 (no symptoms) to 28 (death), assessing motor and sensory functions such as body symmetry, gait, climbing, circling behaviour, forelimb symmetry, compulsive circling, and whisker response, based on previously published scales^29^.

### Organ collection and bacterial culture

6 h, 24 h, 96 h, and 168 h after MCAO mice were euthanized via intraperitoneal injection of a pentobarbital overdose (150 mg/kg), followed by intracardiac perfusion with sterile PBS. Segments of the liver, heart, lungs, spleen, and brainstem were collected, along with blood obtained via cardiac puncture. All samples were homogenized with a tissue grinder in sterile PBS (600 µl) and plated on brain-heart infusion (BHI) agar plates (45 g/l, Biokar), a nutrient-rich medium chosen to support the growth of a broad spectrum of gut-derived bacteria. Plates were incubated in aerobic conditions overnight (O/N) at 37 °C to determine colony-forming units (CFUs). To assess the overall bacterial burden across different organs, we developed a bacterial dissemination index. For each mouse, raw CFU counts from the six mentioned tissues (*i.e.,* liver, lung, spleen, heart, blood, and brainstem) were first converted into a discrete scoring system ranging from 0 to 22. Scores were assigned based on specific intervals: a count of 0 was scored as 0, counts from 1 to 240 were scored as 1, counts from 241 to 480 as 2, and so forth in continuous increments of 240 CFU per score unit. Saturated plates exceeding 5000 CFU were capped at a maximum score of 22. The final dissemination index for each mouse was then calculated by aggregating these individual organ scores.

### Infarct volume assessment

Brains were collected, post-fixed in 4% paraformaldehyde (PFA) for 24 h at 4 °C, and cryoprotected in 30% sucrose for 48 h at 4 °C. Subsequently, brains were frozen in isopentane on dry ice at −25 °C and stored at −80 °C until further processing. Coronal sections (25 µm thick) were obtained using a cryostat. Cresyl violet (CV)-stained sections were prepared by gradual hydration in ethanol, stained with 1 mg/ml CV-acetate in distilled water, gradual dehydration in ethanol, and clearing in xylene^25^. Sections were imaged with a light stereomicroscope at 1.2x magnification (Nikon SMZ25). The infarct volume was determined by multiplying the sum of the infarcted areas on each section (ImageJ Fiji 1.54p software) by the section spacing (700 µm).

### Swiss roll and gut assessment

Colons were isolated, measured, opened longitudinally, and cleaned of faecal content using PBS. The proximal end was inserted into the eye of a 25G needle and gently rolled toward the distal end (so-called Swiss roll preparation^30^). Samples were then post-fixed in 4 % PFA for 24 h at 4°C, and then cryoprotected in 30 % sucrose for 48 h at 4°C. Transversal sections 25 µm thick were obtained using a cryostat. Alcian Blue-stained sections were prepared by hydration through a graded ethanol series, followed by staining with Schiff reagent, rinsing in water, staining with 1% Alcian Blue in 3% acetic acid, another water rinse, and counterstaining with haematoxylin. Sections were then gradually dehydrated through ascending ethanol concentrations and cleared in xylene. Mucin content was evaluated using ImageJ.

### Immunostaining

For immunofluorescence labelling, 25 µm free-floating brain sections and selected colon slices were washed with PBS and blocked and permeabilized with PBS containing 1 % BSA, 0.1 % Triton X-100, and 10 % horse serum for 60 min at room temperature. Sections were then incubated O/N at 4°C with primary antibodies anti-iba1 (1:400, Abcam), anti-peptidoglycan (1:400, Sigma), anti-ZO-1 (1:200, Fisher) in PBS containing 1 % BSA, 0.1 % Triton X-100, and 5 % horse serum. After extensive washing with PBS, sections were incubated with the corresponding secondary Alexa Fluor-conjugated antibodies (1:300, Sigma) and DAPI for nuclear counterstaining in PBS containing 1 % BSA, 0.1 % Triton X-100, and 2 % horse serum for 60 min. For IgG staining, the same protocol was followed using only an Alexa Fluor 594-conjugated goat anti-mouse IgG secondary antibody (Invitrogen A11032). Sections were mounted with Fluor-save (Calbiochem). Images were captured using a Zeiss Z1 slide scanner (Carl Zeiss, Germany) microscope at 20x magnification. Analyses of peptidoglycan, microglia, and ZO-1 were performed with ImageJ. IgG leakage volume was calculated by multiplying the cumulative leakage area per section by the 700 µm inter-section interval, using QuPath software v.0.5.1.

### Flow cytometry

Mesenteric lymph nodes (MLNs) were harvested and processed into single-cell suspensions by mechanical dissociation through a 70-µm nylon cell strainer, followed by enzymatic digestion with collagenase D (0.2 mg/ml) in HBSS supplemented with 10 mM HEPES and 5% FBS. Cell suspensions were subsequently filtered through a 40-µm cell strainer prior to staining. Cells were stained with fixable viability stain 620 (564996, BD Biosciences) according to the manufacturer’s instructions. Nonspecific binding was blocked by incubation with anti-CD16/CD32 antibody (clone 93, 16-0161-85, Invitrogen). Cells were then stained for 20 min at 4 °C with fluorochrome-conjugated anti-mouse antibodies against CD45 (clone 30-F11, BioLegend), CD4 (clone RM4-5, BioLegend), CD8 (clone 53-6.7, BioLegend), and B220 (clone RA3-6B2, BioLegend). Data were acquired on a BD LSR II cytometer (BD Biosciences) and analysed using FlowJo software version 10.9.0.

### Fecal and brain DNA extraction for 16S rRNA gene amplicon sequencing

The DNA was extracted from fecal and brain samples using a standardized protocol involving automated physical disruption via bead beating along with chemical cell lysis using the DNeasy PowerLyzer PowerSoil Kit (Qiagen, Germany) following the manufacturer’s instructions. Mock community DNA was included as positive control for library preparation (Zymobiomics Microbial Community DNA, ZymoResearch, Irvine, CA, USA) and to ensure quality control. Negative controls of DNA extraction and PCR amplification steps were routinely performed in parallel, as well as positive controls, using the same conditions and reagents. Samples were amplified using 16S rRNA V3-V4 regions specific primers (Forward 5′TCGTCGGCAGCGTCAGATGTGTATAAGAGACAGCCTACGGGNGGCWGCAG-3′, Reverse 5′GTCTCGTGGGCTCGGAGATGTGTATAAGAGACAGGACTACHVGGGTATCTAATCC-3′). Cycling conditions were initial denaturation of 3 min at 95 °C followed by 25 cycles of 95 °C for 30 s, 55 °C for 30 s, and 72 °C for 30 s, ending with a final elongation step of 5 min at 72 °C. PCR reactions were purified using AMPure XP beads (Beckman Coulter) with a 0.9× ratio according to manufacturer’s instructions. The primers contain overhangs allowing the addition of full-length Nextera adapters with barcodes for multiplex sequencing in a second PCR step, resulting in sequencing ready libraries with approximately 450 bp insert sizes. Sequencing was performed in an Illumina MiSeq with 2×300 bp reads using v3 chemistry with a loading concentration of 10 pM. In all cases, libraries were spiked with 15% PhiX control to increase the nucleotide diversity of the sequencing run. DNA extraction, library preparation, and sequencing were performed by Microomics S.L. (Barcelona, Spain).

### Bioinformatic sequence analysis

Raw read processing was performed with the QIIME2 pipeline (v2026.4, amplicon distribution)^31^. Adapter sequences were first trimmed with fastp (v1.0.1)^32^. Quality- and length-based filtering were disabled to leverage downstream filtering by DADA2. Read quality was assessed with FastQC (v0.12.1)^33^ / MultiQC (v1.33)^34^, and truncation positions were chosen as the first base at which the mean Phred score fell below Q25. We determined that quality for forward reads did not drop before 300 nt, while for reverse reads the earliest drop was at 225 nt. The DADA2 ^35^ plugin in QIIME2 via qiime dada2 denoise-paired was used to trim and merge the paired-end reads, remove chimeric sequences, and denoise reads for resolving amplicon sequence variants (ASVs). Default parameters were used except for trimming 17 nt and 21 nt from the 5′ ends of the forward and reverse reads (--p-trim-left-f and --p-trim-left-r) to remove the 341F and 805R primers (CCTACGGGNGGCWGCAG and GACTACHVGGGTATCTAATCC, respectively)^36^, and forward and reverse truncating at 295 nt and 225 nt, respectively (--p-trunc-len-f and --p-trunc-len-r). Expected overlap was therefore 56 bp for the ∼464 bp V3–V4 amplicon. Denoising yielded 1,820 ASVs from a total of 896,261 input reads, of which 227,205 (25.3%) were retained as non-chimeric merged reads. The extracted ASVs were taxonomically classified using the sklearn-based Naive Bayes classifier^37^ (scikit-learn v1.7.1, confidence threshold 0.7) trained on the SILVA v138.2 SSURef NR99 database^38^. ASVs not assigned to the domain Bacteria, and any assigned to mitochondria or chloroplasts, were removed from both the feature table and the representative sequences, leaving 1,105 bacterial ASVs and 195,288 reads (86.0% of non-chimeric reads). Next, to conform to up-to-date recommendations^37,39^, filtering based on prevalence and abundance thresholds was implemented. The prevalence filter retained ASVs detected at more than one read in at least 10% of samples, (230 ASVs, 117,441 reads), and the abundance filter retained ASVs whose assigned reads were at least 0.10% of the total reads after the prevalence filter (178 ASVs, 114,119 reads).

Taxonomic abundance and alpha and beta diversity analyses were performed using the Phyloseq R package (v1.56.0)^40^. Rarefaction curves for fecal samples plateaued at around 7,000 reads, which suggested that diversity measures were appropriate. For brain samples, due to the lower number of reads, diversity metrics were not computed. All p-values were adjusted with Benjamini-Hochberg correction. All plots were performed in R v4.5.2 using the R packages ggplot2 v4.0.2^41^ and ggpubr v0.6.3^42^. The SILVA classifier did not originally resolve any ASV to species level, so analyses were performed at family level.

### Statistical analyses

Data are expressed as the mean ± SEM. All experiments were randomized and conducted in a blinded manner. Comparisons between two groups were done using the unpaired Student’s t-test for parametric data or the Mann-Whitney U-test for non-parametric data. For comparisons among multiple groups, one-way ANOVA followed by Fisher’s LSD post-hoc test was used for normally distributed data. For non-parametric datasets, the Kruskal-Wallis test with Dunn’s post-hoc correction was employed. All analyses were performed using GraphPad Prism v8.3.0 software. We considered *p*-values <0.05 as significant.

## Results

### First assessment of spatiotemporal distribution of bacteria after stroke

Given the limited data on bacterial translocation and dissemination after stroke, we first aimed to determine whether this phenomenon occurs in peripheral organs following ischemia. Furthermore, we sought to establish its onset and duration using a mouse MCAO model (Fig. 1A). As controls, we used both naïve and sham-operated mice. In the sham group, the procedure mirrored the MCAO surgery, except that the filament was inserted into the common carotid artery without occluding the middle cerebral artery. This control setup allowed us to rule out bacterial growth resulting from the skin incision, surgical manipulation, filament insertion, or cranial probe placement, confirming that the observed bacterial presence was linked to the ischemic event and not to the surgical procedure.

**Figure 1.**
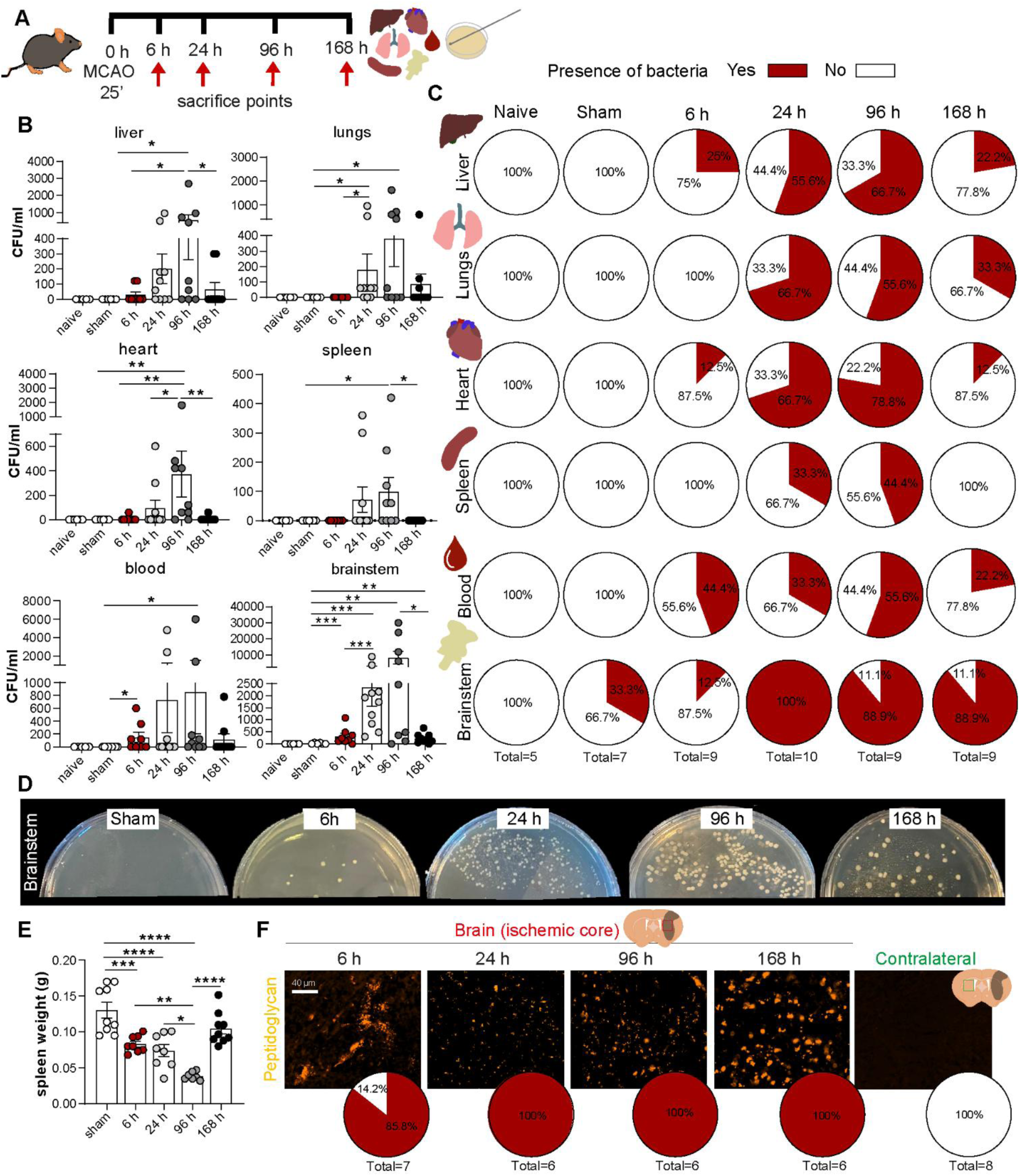
Spatiotemporal distribution of bacteria after stroke. **A.** Experimental design. Mice were subjected to a 25-minute middle cerebral artery occlusion (MCAO) or sham surgery. Animals were sacrificed at 6, 24, 96, or 168 h post-surgery, and liver, lung, heart, spleen, blood, and brainstem samples were collected and cultured on brain-heart infusion (BHI) agar plates**. B.** Bacterial load (CFU/ml) in the indicated organs from naïve, sham, and MCAO groups (n=5 naïve, n=7 sham, n=9-10 MCAO; 3 independent experiments). **C.** Pie charts representing the percentage of mice with an absence (white) or presence (red) of bacteria across the different groups, time points, and organs. **D.** Representative images of brainstem cultures on BHI plates at different time points, illustrating the temporal progression of bacteria. **E.** Quantification of spleen weight across experimental groups as an indicator of immunosuppression (n=8-9; 3 independent experiments). **F.** Representative immunofluorescence images of the ischemic core (striatum, left) and the contralateral healthy hemisphere (right) stained for bacteria (peptidoglycan, yellow) at the different time points. The bottom pie charts represent the percentage of samples positive (red) or negative (white) for bacteria in each group. Data are the mean ± SEM; dots represent individual mice. \**p* < 0.05, \*\**p* < 0.01, \*\*\**p* < 0.001, \*\*\*\**p* < 0.0001.

By culturing multiple organs collected at different time points on BHI agar, chosen for its suitability to culture gut-derived bacteria, we observed a widespread systemic presence of bacteria after stroke. Significant bacterial loads were detected in the liver, lungs, heart, spleen, blood, and brainstem, compared to sham and naïve mice, which showed minimal or no bacteria, respectively (Fig. 1B,C). The minimal bacterial load in sham mice probably reflects bacterial dissemination due to the surgical procedure itself. Interestingly, this dissemination began as early as 6 h post-ischemia (with bacteria detected in 10–45% of samples from the liver, heart, brainstem, and blood, but not in the spleen or lungs). This was followed by a massive bloom at 24 h (affecting 33% to 100% of samples per organ) that reached an observed peak at 96 h (with 44% to 89% of samples testing positive for bacteria), though the actual maximum likely fell between 24 and 96 h for most organs. By 168 h, however, bacterial numbers were significantly reduced, indicating a rapid but transient systemic spread (with 65–100% of samples showing no detectable culturable bacteria by this time point) (Fig. 1B,C).

Remarkably, the brainstem exhibited a particularly pronounced accumulation of bacteria in terms of absolute numbers and prevalence (between 89% and 100% from 6 h onwards) compared to peripheral organs (Fig. 1B-D). This timeline closely coincides with the spatiotemporal systemic immunosuppression (measured by spleen weight) that was evident by 6 h, peaked at 96 h, and almost completely resolved by 168 h (Fig. 1E). Next, we examined the bacterial presence in the brain using an antibody against peptidoglycan, a major constituent of the bacterial cell wall, allowing the detection of intact bacteria or bacterial fragments within the tissue. This analysis revealed that bacteria accumulated exclusively within the ischemic core (Fig. 1F). Importantly, no bacteria were detected in perilesional or unaffected areas, the contralateral healthy hemisphere, or in naïve and sham mice, confirming that the observed signal was genuine and not due to contamination (Fig. 1F, Supplementary Fig. 2). Spatiotemporal analysis demonstrated for the first time that bacteria infiltrated the brain as early as 6 h post-ischemia and persisted at 24 h, 96 h, and 168 h (observed in 100% of evaluated brains from 6 h onwards). This prolonged cerebral colonization contrasts sharply with that of peripheral organs, where bacterial loads declined by 168 h (Fig. 1B-D).

### Analysis of brain bacteria and gut dysbiosis and their association with post-stroke functional outcomes and survival

To confirm whether the bacteria detected in the brain originated from the gastrointestinal tract, we performed 16S rRNA gene sequencing on paired brain and stool samples collected at day 4 post-surgery (Fig. 2A). This time point was selected to coincide with the peak of systemic bacterial dissemination and profound immunosuppression (Fig. 1). To ensure rigorous quality control and account for potential environmental contamination, we included sham-operated controls and applied abundance and prevalence filters in the bioinformatic processing. Given the inherently low bacterial biomass of the brain, applying these strict controls is imperative, as standard 16S rRNA gene amplicon sequencing without rigorous background noise filtering can easily yield misleading false positives. No bacterial sequences were detected in the brains of any sham-operated mice (n=3). In contrast, two out of four MCAO brain samples harboured bacterial sequences. In one of these samples (MCAO 1), a minor fraction of the sequences was assigned to the family *Erysipelotrichaceae* (*i.e.,* a common gut symbiont), supporting the hypothesis of gut-to-brain bacterial translocation (Suppl. Fig. 3A). The vast majority of the remaining sequences in this sample, as well as all sequences detected in a second sample (MCAO 3), mapped to the bacterial kingdom but could not be resolved at lower taxonomic ranks, likely due to limited 16S resolution and low sequencing depth.

**Figure 2:**
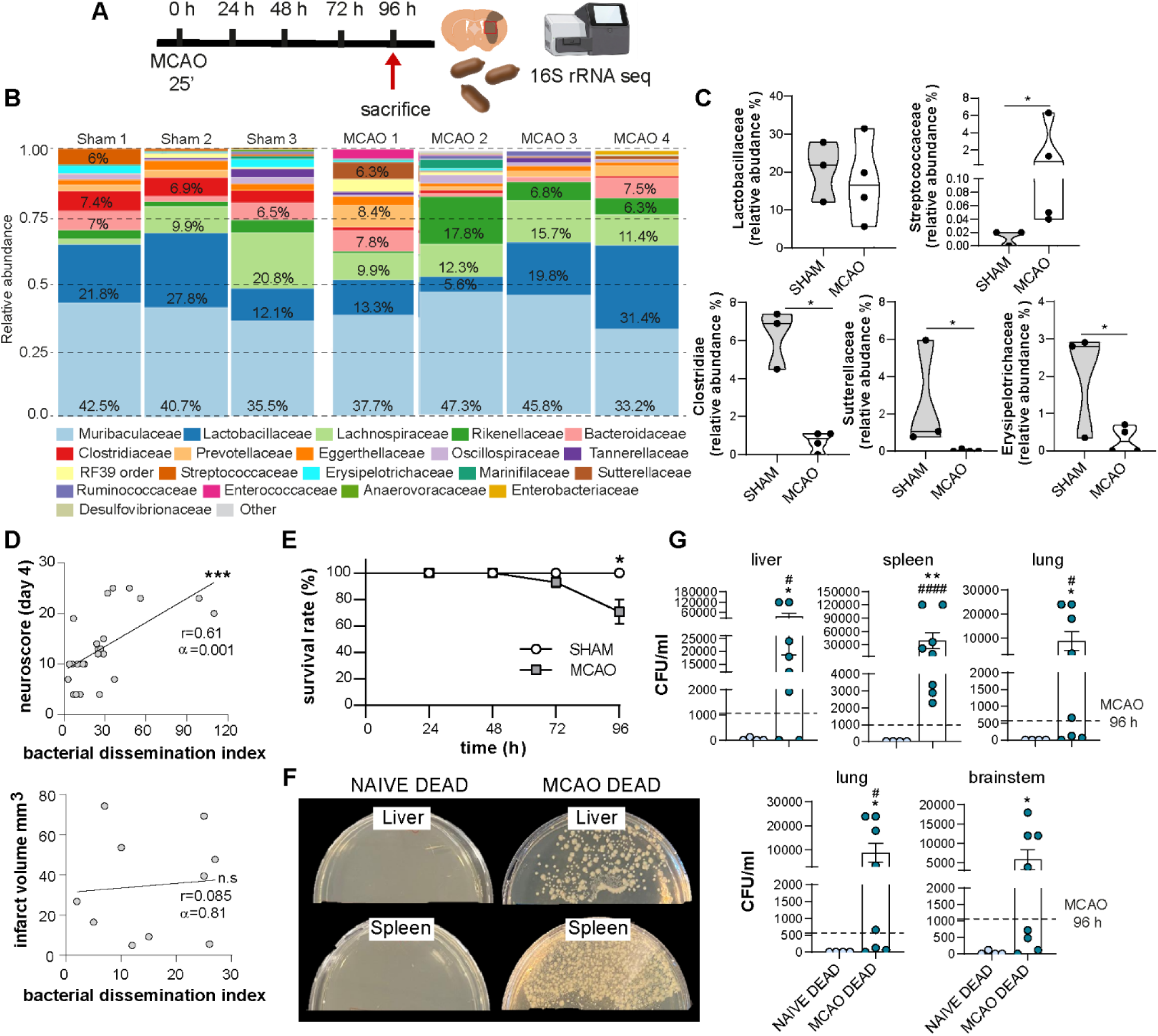
Post-stroke gut dysbiosis and correlation with functional outcomes and survival. **A.** Experimental design. Mice were subjected to transient middle cerebral artery occlusion (MCAO; 25 min) or sham surgery and sacrificed at 96 hours post-reperfusion for sample collection (brain and stool) and 16S rRNA gene sequencing. **B.** Stacked bar plots illustrating the relative abundance of bacterial families in the stool microbiome of sham (n=3) and MCAO (n=4) mice at day 4**. C.** Violin plots comparing the relative abundance of specific bacterial families (*Lactobacillaceae, Streptococcaceae, Clostridiaceae, Sutterellaceae,* and *Erysipelotrichaceae*) between sham and MCAO groups. **D.** Pearson correlation analysis evaluating the relationship between the Bacterial Dissemination Index (BDI, a composite of CFUs across the different organs) and neuroscore at day 4 (top graph, N=25) as well as infarct volume (bottom graph, N=10). **E.** Survival curves for sham and MCAO mice over the 96-hour observation period. **F.** Representative images of BHI agar plates cultured with liver and spleen homogenates from overnight-intact deceased naïve mice and non-surviving MCAO mice. **G.** Quantification of bacterial loads (CFU/ml) in liver, spleen, lung, heart, and brainstem in deceased MCAO mice vs. euthanized naïve controls subjected to an overnight post-mortem interval before tissue collection and surviving MCAO mice sacrificed at 96 h (dashed line). Data are the mean ± SEM; dots represent individual mice. *^/#^*p*<0.05, **^/##^*p*<0.01, ***^/###^*p*<0.001, ****^/####^*p*<0.0001; asterisks (*) indicate significant differences relative to the MCAO group. In G, asterisks (*) indicate significant differences relative to the naïve dead group, and hash signs (#) denote significance compared with MCAO CFUs at 4 days (dashed line).

Analysis of the corresponding stool samples revealed that, despite no significant alterations in overall alpha or beta diversity between the sham and MCAO groups (Suppl. Fig. 3B,C), there were marked shifts in the relative abundance of specific taxa (Fig. 2B,C). While predominant families such as *Lactobacillaceae* remained stable, indicating that the post-stroke dysbiosis was highly targeted, the relative abundance of *Streptococcaceae*, a bacterial family commonly associated with PSIs, was significantly elevated in MCAO mice (Fig. 2B,C). Conversely, the relative abundance of potentially protective taxa, including *Sutterellaceae* and the SCFA-producing *Clostridiaceae*, was reduced. Notably, *Erysipelotrichaceae*, the same taxon detected in the brain, exhibited a decreased relative abundance in the stool of MCAO mice (Fig. 2B,C), further supporting the potential occurrence of gut-to-brain translocation.

Interestingly, at this time point (4 days), higher neuroscore, indicative of more severe sensory and motor deficits, positively correlated with an increased Bacterial Dissemination Index (BDI), calculated as a composite of the CFUs detected across various organs (see Methods). In contrast, ischemic volume did not correlate with a higher BDI (Fig. 2D). This dissociation might indicate that the stroke-induced systemic response driving bacterial translocation is reflected in the functional outcome of the animal, but remains independent of the extent of local brain damage. Furthermore, approximately 75% of non-surviving MCAO mice (succumbing primarily on days 3 or 4; Fig. 2E) displayed a massive systemic bacterial burden compared to surviving MCAO mice evaluated at day 4 (dashed line; Fig. 2F,G). To ensure that bacterial loads in mice that died were not merely artifacts of post-mortem overgrowth, healthy naïve mice were euthanized and kept at room temperature overnight to simulate the post-mortem interval. Subsequently, their organs were harvested and cultured. The complete absence of bacterial growth in these BHI cultures confirmed that the extensive bacterial spread was driven by the ischemic pathology rather than post-mortem artifacts (Fig. 2G). However, stroke-induced mortality was not exclusively infection-driven, as approximately 25% of the deceased mice showed no detectable bacterial growth.

### Post-stroke colon dysfunction and gut barrier integrity are modulated by SCFAs

Considering the lack of effective prophylaxis for PSIs, the well-documented benefits of SCFAs in other models^18–20,22,23,43–45^, and the significant post-ischemic depletion of SCFA-producing taxa demonstrated by our data (Fig. 2C) and previous studies (reviewed in ^2^), we investigated the potential therapeutic administration of SCFAs in our model. We first tested whether administering an SCFA mixture (acetate, butyrate, and propionate; delivered every 24 h starting one day post-stroke) could confer protection against systemic bacterial dissemination (Fig. 3A). SCFA treatment resulted in a significant reduction in bacterial load in the liver, lungs, and spleen, with an especially pronounced decrease observed in the brainstem. No significant differences were found in the heart or blood (Fig. 3B,C).

**Figure 3.**
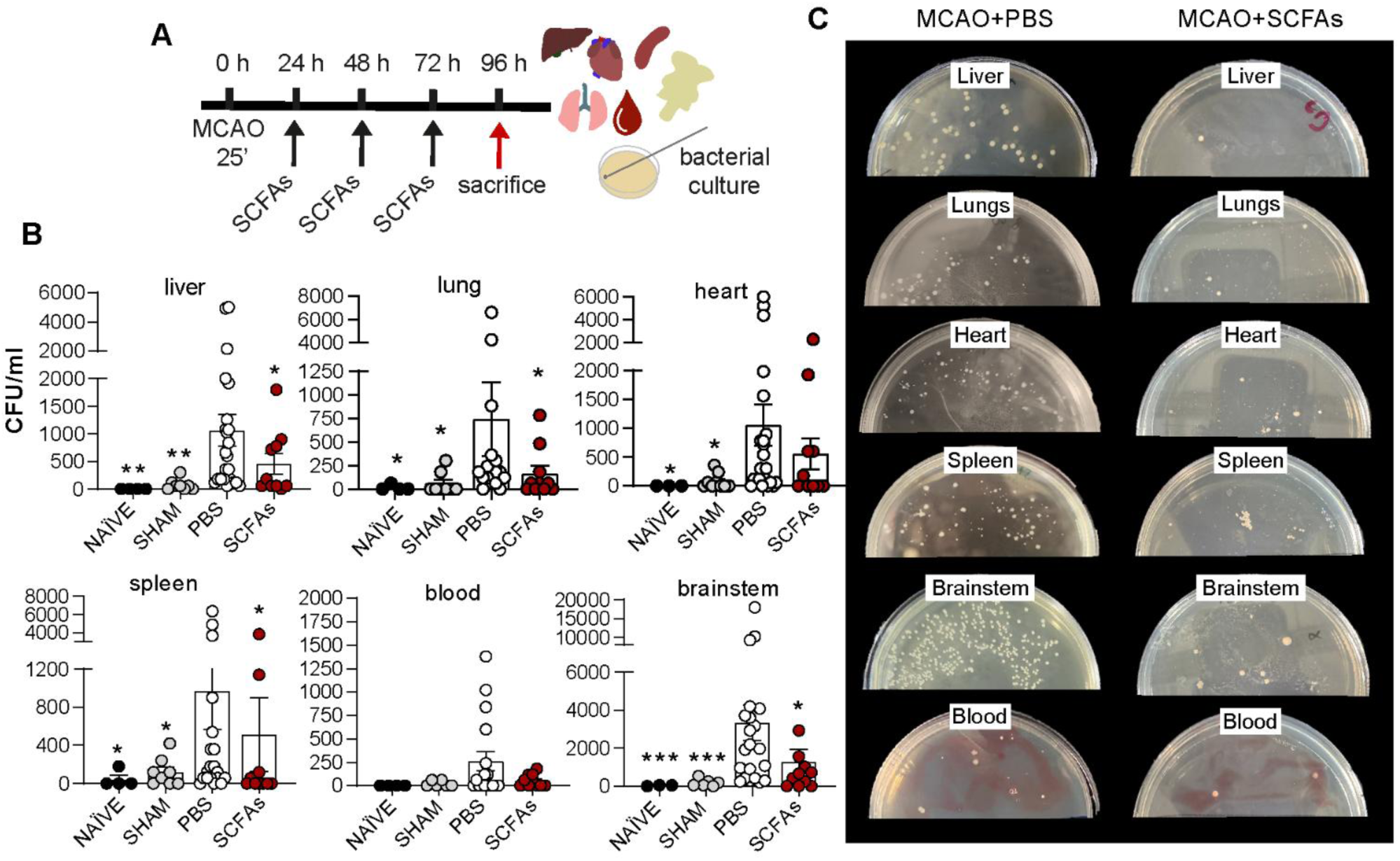
SCFAs reduce bacterial burden after stroke. **A.** Experimental design. Mice were subjected to 25 min MCAO or *sham* surgery. MCAO mice received daily intraperitoneal injections of either PBS or an SCFA mixture (acetate:butyrate:propionate, 60:20:20, 6 mmol/kg), starting one day post-stroke. Animals were sacrificed on day 4, and the liver, lung, heart, blood, spleen, and brainstem were collected for culture on BHI agar plates. **B.** Bacterial load (CFU/ml) in the mentioned organs from naïve, *sham*, and MCAO groups (naïve N=3–4; *sham* N=6–9; MCAO+PBS N=17–23; MCAO+SCFAs N=10–11; 5 independent experiments). **C.** Representative images of BHI agar plates from MCAO+PBS and MCAO+SCFAs mice. Data are the mean ± SEM; dots represent individual mice. \**p*<0.05, \*\**p*<0.01, \*\*\**p*<0.001; asterisks (*) indicate significant differences relative to the MCAO + PBS group.

To understand the mechanisms behind the modulation of bacterial translocation by SCFAs, we next evaluated the effects on gut barrier integrity (Fig. 4A), since maintaining the integrity of the gut epithelium is critical for preventing the systemic dissemination of commensal bacteria. SCFA treatment attenuated the post-stroke reduction in colon length, a common marker of intestinal dysfunction, observed in MCAO mice (Fig. 4B). SCFA-treated mice also attenuated the reduction in faecal pellet output induced by MCAO, particularly on days 3 and 4 post-stroke (Fig. 4B), suggesting that these metabolites may counteract stroke-induced dysmotility. Histological analysis of the colon using the Swiss-roll technique revealed a post-stroke reduction in the immunoreactivity of the tight-junction protein ZO-1 throughout the epithelium. Notably, this loss of epithelial integrity was partially restored following SCFA treatment (Fig. 4C,D). Furthermore, the density of Alcian Blue-positive mucins, which act as a critical physical barrier restricting luminal bacteria and preventing their translocation across the intestinal wall, was drastically reduced after MCAO, but significantly recovered following SCFA administration (Fig. 4C,D).

**Figure 4.**
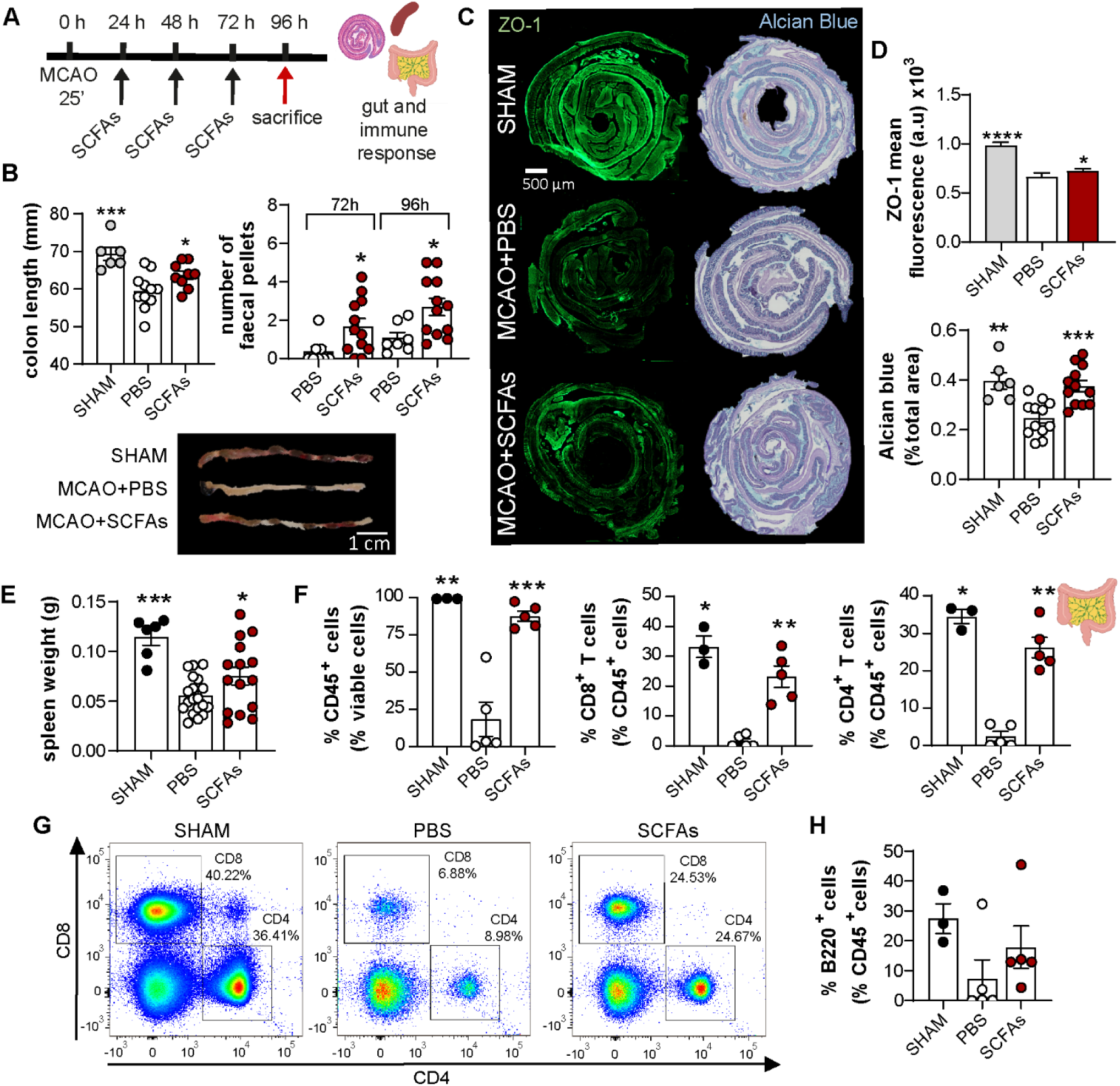
SCFAs preserve colon length and gut barrier integrity and modulate immune response. **A.** Experimental design. Mice were subjected to 25 min MCAO or *sham* surgery. MCAO mice received daily intraperitoneal injections of either PBS or an SCFA mixture (acetate:butyrate:propionate, 60:20:20, 6 mmol/kg), starting one day post-stroke. Animals were sacrificed on day 4. **B.** Top. Quantification of colon length and fecal pellet in SHAM, MCAO+PBS, and MCAO+SCFAs groups (N=6–11; 4 independent experiments). Bottom. Representative colon images. **C.** Representative colon sections (Swiss rolls) immunostained for ZO-1 (green) to label tight junctions, and stained with haematoxylin and Alcian Blue to visualise mucins (blue). **D.** Top, ZO-1 fluorescence intensity quantification (N=3-5, 10 ROIs per colon; 2 independent experiments). Bottom, Alcian Blue quantification expressed as percentage of total area (N=6-12; 2 independent experiments). **E.** Quantification of spleen weight across experimental groups (N=6-19, 5 independent experiments). **F.** Flow cytometry analysis of MLNs: percentage of total CD45^+^ hematopoietic cells (left, gated on viable cells), CD8^+^ T cells (middle, gated on CD45^+^ cells), and CD4^+^ T cells (right, gated on CD45^+^ cells) (N=3-5, 2 independent experiments). **G.** Representative flow cytometry plots illustrating CD4^+^ and CD8^+^ T cell populations across groups. **H.** Percentage of total B220^+^ cells (gated on CD45^+^ cells). Data are the mean ± SEM; dots represent individual mice. \**p*<0.05, \*\**p*<0.01, \*\*\**p*<0.001, \*\*\*\**p*<0.0001; asterisks (*) indicate significant differences relative to the MCAO + PBS group.

As bacterial dissemination is also associated with profound immunosuppression, and given the immunomodulatory properties of SCFAs^24^, we evaluated the impact of SCFAs on the systemic immune response to determine whether the reduction in bacterial translocation was linked to immune modulation. Notably, SCFA treatment partially reversed the marked stroke-induced immunosuppressive state, as evidenced by a significant recovery in spleen weight (Fig. 4E). Given that MLNs are the primary draining lymph nodes of the intestine and critical sites for immune priming in response to gut-derived antigens^46^, we next examined immune cell populations within the MLNs via flow cytometry. We observed a significant depletion of total hematopoietic cells (CD45^+^) in MCAO mice, which was significantly restored following SCFA administration. Within the CD45⁺ compartment, the percentages of CD4^+^ and CD8^+^ T cells that were heavily reduced post-stroke were nearly completely recovered in the SCFA-treated group (Fig. 4F,G), while the B-cell population remained less affected by stroke and SCFA treatment (Fig. 4H).

### SCFA treatment attenuates brain injury and cerebral bacterial translocation after stroke

Given that SCFA administration reduced bacterial dissemination, protected gut integrity, and modulated the immune response, we finally examined whether it might also mitigate the ischemic lesion and improve associated outcomes (Fig. 5A). Although no significant difference was observed in the 28-point neuroscore between saline- and SCFA-treated mice, animals treated with SCFAs displayed a higher survival rate compared with PBS-treated controls at day 4 (Fig. 5B). Moreover, SCFA-treated mice exhibited a significant reduction in the total infarct volume on day 4, which was characterized by marked decreases in both striatal and hippocampal lesions (Fig. 5C). SCFAs also ameliorated post-stroke BBB breakdown, as evidenced by a significant reduction in IgG leakage from the blood into the brain parenchyma (Fig. 5D).

**Figure 5.**
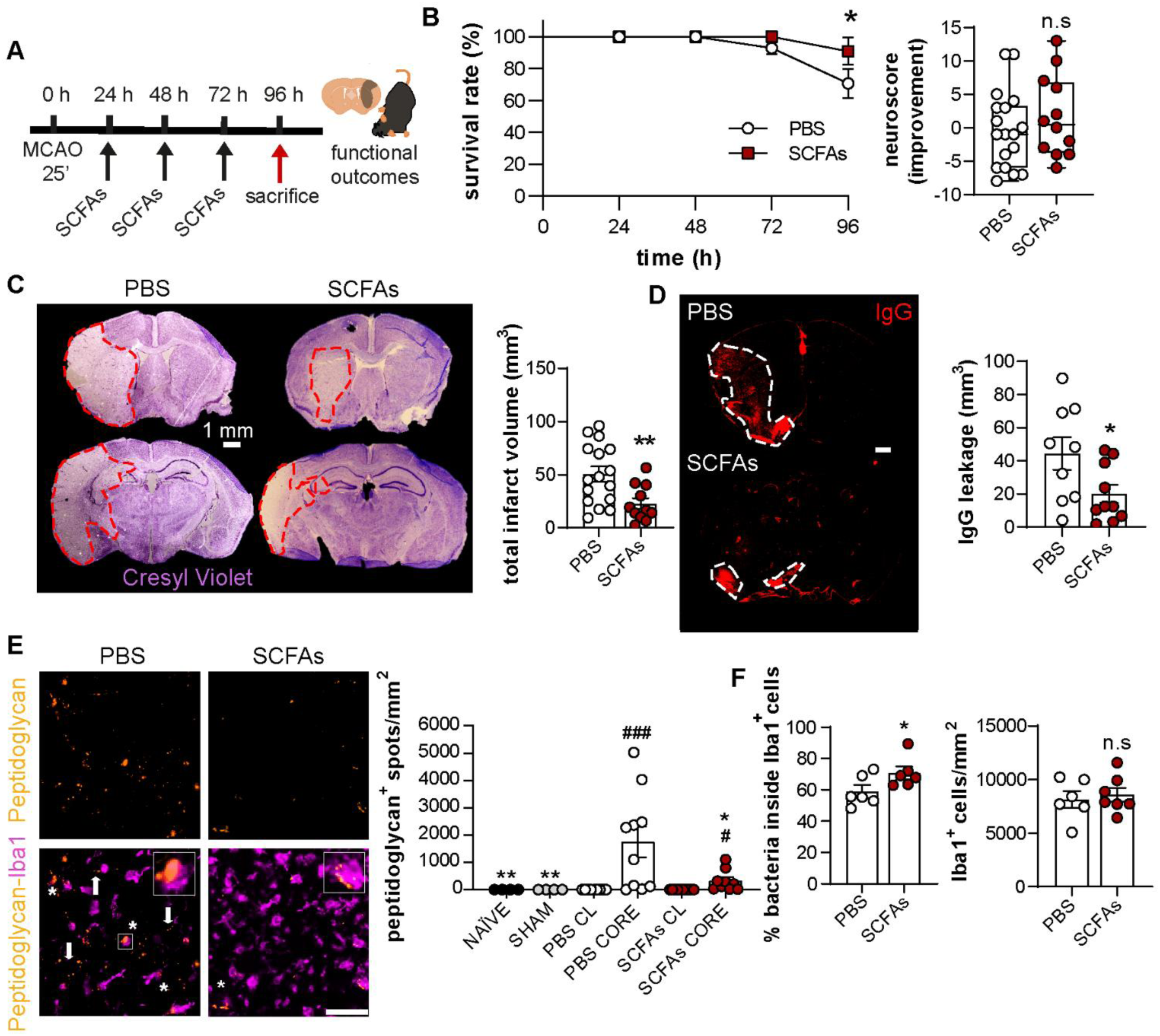
SCFAs reduce brain injury, BBB leakage, and bacterial presence in the brain. **A.** Mice subjected to MCAO received daily treatment with either PBS or an SCFA mixture (acetate:butyrate:propionate, 60:20:20; 6 mmol/kg), starting one day post-stroke, and were sacrificed on day 4 for comparison with *sham* mice. **B.** Percentage of survival over the course of the experiment (left) and neuroscore improvement between day 1 and day 4 (positive values indicate improvement) on a 28-point scale (right) (N=12–21; 5 independent experiments). **C.** Left, representative images of brain lesions in the striatum and hippocampus (red dashed lines) stained with cresyl violet. Scale bar: 1 mm. Right, infarct volume quantification in the whole brain (N=11–17; 3 independent experiments). **D.** Representative images of BBB leakage (IgG, red; delineated with white dashed lines) and IgG leakage volume quantification (N=9–10; 3 independent experiments). Scale bar, 1 mm. **E.** Left, images of the ischemic core (striatum) stained for bacteria (peptidoglycan, yellow) and Iba1 (microglia, magenta). Asterisks indicate bacteria being phagocytosed by microglia/macrophage-like cells (high-magnification insets) and arrows indicate bacteria within the brain parenchyma. Right, quantification of peptidoglycan spots/area in naïve, *sham*, MCAO+PBS, and MCAO+SCFAs animals (comparing the ischemic core to the contralateral non-injured side, CL) (N=4–9; 3 independent experiments). Scale bar: 20 µm; insets: 3x magnification. **F.** Quantification of bacteria within Iba1^+^ cells (%) and Iba1^+^ cells density (cells/area) in the ischemic core (striatum) (N=6, 2–3 ROIs per mouse, 2 independent experiments). Data are presented as mean ± SEM; dots represent individual mice. *^/#^*p*<0.05, \*\**p*<0.01, ^###^*p*<0.001. Asterisks (*) indicate significant differences relative to the MCAO + PBS group and hash signs (#) denote significance compared with the CL side.

Importantly, SCFA treatment markedly reduced the peptidoglycan positive signal, reflecting reduced presence of bacteria or bacterial fragments in the ischemic core (Fig. 5E). Notably, these bacteria or bacterial fragments were partially phagocytosed by Iba1-positive microglia/macrophage-like cells (Fig. 5E, F). Interestingly, we observed that a higher proportion of the detected peptidoglycan signal was localized intracellularly within microglia/macrophage-like cells in the SCFA-treated group compared to PBS controls, independent of Iba1+ cell density, which remained comparable between groups (Fig. 5F).

Taken together, these data provide the first insights into the spatiotemporal dynamics of bacterial dissemination following stroke, revealing a peripheral burden that emerges and spreads rapidly before dissipating relatively early, in stark contrast to the persistent colonization observed in the central nervous system. Furthermore, we demonstrate that SCFAs restore intestinal barrier integrity and reverse stroke-induced immunosuppression, replenishing CD4+ and CD8+ T-cell populations in the MLNs. These systemic effects, potentially reinforced by direct antimicrobial properties or hitherto undefined pathways, collectively limit bacterial translocation to both peripheral organs and the brain. Importantly, SCFA administration also provides significant neuroprotection by reducing infarct volume and preserving BBB integrity. While the attenuation of bacterial burden and these neuroprotective actions may stem from distinct mechanisms, their convergence likely drives the improved survival observed in our model, highlighting SCFAs as a promising multimodal therapeutic strategy against stroke and PSIs.

## Discussion

Stroke is increasingly recognized as a systemic disorder rather than merely a brain-centred event. It involves not only brain injury, neuroinflammation, and BBB disruption, but also complex local and systemic immune responses, along with disturbances in the brain-immune-gut axis^2^. Within the intricate spatiotemporal dynamics of stroke and its outcome, PSIs remain among the most life-threatening complications^4^. Despite the fact that the phenomenon of bacterial translocation has been extensively discussed since its first formal description in the 60’s by Wolochow *et al.*^47^, research has since largely centred on conditions associated with direct injury to the gut barrier, leading to its disruption (*e.g.,* inflammatory bowel disease, colorectal cancer), with comparatively limited attention to brain disorders. It is only very recently, and initially met with scepticism, that pioneering investigation of bacterial translocation in neurodegenerative disorders^48^ and stroke^5,6,49^ has begun. Following the original study reporting that most bacteria detected in patients with PSIs are intestinal commensals that translocate after stroke^6^, our findings confirmed the presence of bacteria in multiple organs after MCAO, including the lungs, liver, heart, and spleen, as has also been observed by others^5,6,50^. Importantly, this phenomenon was linked to the ischemic process, as it differed significantly from *sham*-operated animals, ruling out the surgical procedure as a source of infection. This link is further supported by evidence from different stroke models in germ-free mice, which show that infections do not occur in the absence of a commensal microbiota^6,49^. Interestingly, and in line with these reports, we also observed that not all mice subjected to stroke exhibited culturable bacteria. This finding is consistent with the inherent heterogeneity of stroke outcomes, which persists even within highly standardized experimental models.

To our knowledge, this is the first report to assess the spatiotemporal distribution of bacteria after stroke in both the periphery and central nervous system. Other reports have evaluated bacterial dissemination post-stroke at 24 h^5,6,49^, but not at earlier or later time points. The striking detection of viable bacteria in peripheral organs as early as 6 h suggests that gut barrier failure occurs rapidly. This massive bacterial bloom, which peaked at approximately 96 h, closely aligned with the transient window of stroke-induced immunosuppression, as evidenced by maximal spleen atrophy. The subsequent reduction in peripheral bacterial loads by 168 h indicates that once systemic immunity—and likely the intestinal barrier—begins to recover, peripheral organs successfully regain their antimicrobial clearance capacity, effectively resolving systemic dissemination.

In stark contrast to this peripheral clearance, our data reveal a markedly different dynamic within the central nervous system. The brainstem exhibited an exceptionally rapid and pronounced microbial accumulation that persisted for at least 168 hours, suggesting a prolonged colonization beyond our observation window. Similarly, the ischemic core demonstrated sustained bacterial component persistence throughout the entire study period without signs of clearance. Although peripheral immune cells actively infiltrate the infarcted tissue, their relative accumulation is lower compared to peripheral organs. Furthermore, the profoundly hostile microenvironment—characterized by hypoxia, necrotic debris, and impaired perfusion—likely compromises the antimicrobial efficacy of both these infiltrating cells and resident microglia. Confronted with a sheer volume of cell debris, these immune populations may become functionally exhausted, failing to effectively phagocytose and clear the invading bacteria. Ultimately, this chronic persistence of bacteria/bacterial fragments within the brain parenchyma could act as a continuous inflammatory trigger. This persistence may cause profound secondary damage, acting not only through potential dissemination but also by inducing microglial priming and exacerbating the neuroinflammatory response. These factors could critically compromise long-term recovery and warrant further investigation.

Consistent with our findings, bacterial presence in the brain has been reported in other neurological conditions, such as Alzheimer’s disease^51–54^. However, these observations remain a subject of ongoing debate, since it is unknown whether these microbes are active drivers of neuroinflammation and amyloid deposition, or are merely opportunistic colonizers crossing a compromised BBB. Furthermore, the detection of low-level bacterial signatures in healthy controls raises questions about whether these findings represent real infections or simply environmental contamination^51–54^. Interestingly, oral microbiota has been identified within the thrombi of stroke patients, with the presence of specific oral species correlating with increased mortality^55^.

In this sense, our findings provide novel evidence of bacterial presence in the brain after stroke, corroborating and extending recent observations by Peh *et al*.^49^. We demonstrated that this bacterial accumulation is strictly confined to the ischemic core, remaining undetectable in perilesional areas or the contralateral hemisphere. Although the morphological diversity and size range of the peptidoglycan-positive signals suggest a polymicrobial presence, standard imaging cannot definitively distinguish between viable bacteria, dormant cells, or residual bacterial cell wall fragments. To characterize this bacterial presence, we performed 16S rRNA gene sequencing, which identified sequences assigned to the gut commensal family *Erysipelotrichaceae* within the ischemic brain. Although the absolute read count for these sequences was relatively low, this is an expected outcome when applying 16S rRNA gene sequencing to low-biomass environments. In accordance with current best practices for microbiome research^39,56^, we applied rigorous prevalence and abundance filters to confidently exclude contamination. Furthermore, our methodology incorporates robust controls (*e.g.,* sham-operated mice confirmed to lack peptidoglycan) that are typically absent in the field. Moreover, the sequencing depth and quality achieved here represent a substantial improvement over the only previous comparable attempt in this area^49^ (which lacked both appropriate controls and abundance/prevalence filtering). For instance, while this previous work reported a median of 294 paired-end reads per brain sample, our post-filtering data yielded a median of 81,829 reads. Coupled with superior sequencing quality, these methodological strengths ensure that the detected signals represent genuine biological presence rather than sequencing artifacts. While our comprehensive approach provides strong evidence of bacterial presence, we explicitly acknowledge the inherent limitations of our current study regarding sample size and taxonomic resolution. To overcome these constraints, future studies should employ advanced approaches—such as deep shotgun metagenomics or full-length 16S rRNA sequencing^57,58^—coupled with larger sample sizes and multi-organ experimental designs to definitively identify these translocated pathobionts.

Crucially, our study also provides the first evidence of bacterial presence in the brainstem following stroke. The high bacterial burden observed in this region suggests that translocation may occur via non-canonical pathways, such as the vagus nerve, a mechanism also proposed in other neurological disorders^48^. As a direct anatomical bridge between the gastrointestinal tract and many organs as the lungs, heart, or brainstem^59^, the vagus nerve could account for the low bacterial loads detected in the bloodstream, both in our study and in previous reports^6,49^. Indeed, some evidence suggests that bacterial translocation is reduced following vagotomy or vagal blockade^48,49,60^, though the precise mechanisms underlying gut-to-brain dissemination remain incompletely understood. It is highly plausible that a dual neural and hematogenous mechanism is also at play. Because the bloodstream acts as a rapid transit conduit, circulating bacteria may be swiftly cleared and seeded into target organs, which could also explain transient or low bacterial detection in blood compartments. Furthermore, the remarkably rapid systemic dissemination observed as early as 6 hours post-ischemia temporally coincides with the initial disruption of the BBB^61^. This rapid timeline, coupled with the exclusive localization of bacteria within the ischemic core (*i.e.,* where the BBB is disrupted), strongly points to the fact that these microbes might also infiltrate the brain parenchyma through the compromised local vasculature.

Our findings revealed a highly targeted stroke-induced intestinal dysbiosis, characterized not by a global collapse of microbial diversity (*i.e.,* no differences in alpha or beta diversity, and no changes in major taxa like *Lactobacillaceae*), but by profound functional shifts within specific bacterial families. The significant expansion of *Streptococcaceae* might be particularly detrimental in this context. As opportunistic pathobionts, these bacteria are known to produce virulence factors and enzymes^62^ that can degrade mucosal defensive barriers, which might drive local tissue damage. In the context of stroke-induced immunosuppression, their overgrowth could exacerbate systemic inflammation and might serve as a primary bacterial reservoir for severe PSIs, such as pneumonia. Concurrently, the marked depletion of *Clostridiaceae* might severely compromise intestinal structural integrity. Because this family is a major producer of butyrate, a SCFA that acts as the primary energy source for colonocytes and upregulates the expression of tight junction proteins (*e.g.,* claudins and occludin)^63^, the loss of these keystone SCFA producers may directly dismantle epithelial homeostasis, potentially precipitating the leaky gut phenotype. This barrier dysfunction might be further compounded by the reduction of *Sutterellaceae*, commensals known for their crucial immunomodulatory functions^64,65^. By regulating mucosal immune tolerance, their depletion could remove essential checks on local inflammatory responses, fostering a pro-inflammatory environment highly conducive to bacterial invasion. Most intriguingly, the concomitant reduction of *Erysipelotrichaceae* in the fecal microbiome and its detection within the ischemic brain core suggests a direct gut-to-brain translocation axis. Known for possessing highly immunogenic surface antigens^66^, the decline of this taxon in the stool might not merely reflect luminal death. Rather, it might indicate their physical migration across the compromised intestinal epithelium, disseminating systemically before colonizing the infarcted brain tissue, where their immunogenic nature could further exacerbate local neuroinflammation. Ultimately, the specific bacterial species that colonize different organs, and whether these populations differ by site, as well as the underlying mechanisms driving their systemic dissemination, warrant further investigation and will be the focus of future studies. Moreover, it is critical to acknowledge that our culture findings are constrained by the ‘iceberg paradigm’ of microbiology^67^, since the majority of bacteria remain currently unculturable, the identified bacteria represent only a fraction of the microbiota capable of growing under specific aerobic laboratory conditions.

Currently, no prophylactic strategies are available against PSIs, as antibiotic prophylaxis has not demonstrated improvement in stroke mortality or functional outcomes^7^. Furthermore, in view of the increasing challenge of antibiotic resistance, the development of alternative therapeutic approaches is of critical importance. In this context, the modulation of PSIs through microbial-derived metabolites represents a promising and innovative strategy. Our results demonstrated for the first time that SCFAs can modulate bacterial translocation in stroke, likely through multiple, yet-to-be-defined mechanisms. One such mechanism may involve the preservation of gut barrier integrity (particularly through maintenance of tight junctions and mucin layers), which might limit bacterial dissemination to peripheral organs. Notably, the protective effects of SCFAs on the gut barrier have been reported in other disease contexts (reviewed in ^68^). For example, butyrate is known to regulate the assembly of tight junctions and protect the intestinal barrier^21^. In our study, SCFA administration preserved colon length and increased faecal output, suggesting a significant restoration of gastrointestinal motility. Such effects have been previously described in other gastrointestinal disease models (reviewed in ^69^), where SCFAs influence gut motility by modulating enteric neural activity, neurotransmitter release, calcium signalling, and smooth-muscle contractility^70^. Nevertheless, we cannot exclude the possibility that increased dietary intake in the treated group may have acted as a confounding factor. Given that food consumption was not strictly quantified, it remains possible that the observed benefits on colon length and faecal output were partially mediated by an enhanced nutritional recovery following SCFA administration, as suggested by others^69^.

Furthermore, by counteracting post-stroke immunosuppression, SCFAs may restore the immune response and facilitate bacterial clearance across multiple organs, likely through T-cell modulation. Our findings specifically highlight post-stroke restoration of T cell populations within the MLNs, which serve as the primary site for the induction of mucosal immune responses^46^. Given that MLNs act as a critical checkpoint between the colon and the systemic circulation^71^, the SCFA-mediated modulation observed in this compartment suggests a reinforcement of the intestinal immune barrier and may promote a more tolerogenic environment, potentially limiting pathogen egress while maintaining immune homeostasis. However, the precise relationship between these T-cell shifts in the MLNs and their subsequent recruitment to the colonic lamina propria remains to be fully elucidated. Future investigations should focus on characterizing the spatiotemporal trafficking of these cells to different organs and identifying the specific functional subsets involved (*e.g.,* Tregs or Th17 cells) to better understand how MLN-primed T-cells contribute to long-term gut and brain recovery. Additionally, we cannot rule out the possibility that SCFAs also exert direct antimicrobial effects, either by inhibiting microbial growth or attenuating bacterial virulence, as has been extensively documented^24^.

Beyond gut barrier and immune modulation, SCFAs might prevent bacterial entry to the brain by protecting the BBB. Consistent with this, germ-free mice exhibit reduced expression of key tight junction proteins, resulting in increased BBB permeability from fetal development through adulthood^72^. Notably, the recolonization of these mice with SCFA-producing bacterial strains restores BBB integrity^72^. These findings are further corroborated by *in vitro* evidence, demonstrating that propionate treatment attenuates lipopolysaccharide-induced disruption in brain endothelial cells^73^. Moreover, SCFAs may also modulate the glial and local inflammatory response in the brain, in line with omics-based studies that suggest that microglia are the primary brain targets of SCFAs following stroke^23^. We observed that, despite no significant difference in Iba1+ cells density between treatments, there was an increased number of peptidoglycan-positive spots within the microglia/macrophage-like cells of SCFA-treated mice. This suggests that SCFAs may simultaneously prime or recruit these cells toward a more efficient phagocytic clearance of pathogens, effectively containing the infection at the site of infiltration. In this context, antibiotic-induced perturbations in gut microbial diversity lead to altered microglial morphology and impaired neuroinflammatory signalling (reviewed in ^8^). Additionally, both *in vivo* and *in vitro* evidence indicate that butyrate induces a shift toward a homeostatic profile, inhibiting LPS-induced inflammation^74^. Nevertheless, as our study provides a snapshot at a single time point, we cannot fully characterize the spatiotemporal dynamics of the microglial response, aspects that remain to be fully mapped in future longitudinal studies.

Since it has been previously demonstrated that SCFAs can cross the BBB^75^, a direct effect on neuronal survival or regeneration remains a plausible mechanism. While beyond the scope of the present study, others have reported that SCFAs modulate neurotransmitter levels and neurotrophic factor expression, thereby influencing synaptic plasticity and neuronal connectivity^8,9^. Whether the neuroprotective benefits observed here stem from direct enhancement of neuronal survival or from a concerted regulation of neuroinflammation, BBB integrity, and systemic immune responses remains to be fully elucidated. Regarding the neuroprotective effects of SCFAs, our study is, to our knowledge, the first to report a direct reduction in infarct volume in the MCAO model after post-acute administration. Previous studies employing other stroke models, such as the photothrombotic model, or alternative SCFA administration routes (*e.g.,* drinking water) have reported improvements in functional stroke outcomes or modulated glial/immune responses without a corresponding reduction in lesion size^21–23^. In contrast, our findings demonstrate that SCFA administration, even when initiated at a post-acute time point (24 h), significantly reduces brain injury. This result is particularly relevant given the narrow therapeutic window of current stroke interventions for intravenous thrombolysis and endovascular thrombectomy (up to 24 h after symptom onset). The fact that this treatment effectively mitigates cerebral damage while simultaneously limiting bacterial translocation and preserving the intestinal barrier underscores the potential of SCFAs as a multimodal and translationally viable therapeutic strategy. Interestingly, we did not detect significant differences in behavioural outcome after stroke despite the improved survival, which may be attributable to the assessment scale used. Notably, to date, no behavioural or scoring models have been specifically developed to evaluate PSIs, as the associated behavioural changes are often very subtle. Signs such as piloerection, body weight loss, fever, or shivering are commonly included in MCAO scoring systems, but it is unclear whether these manifestations reflect the neurological lesion itself or have been incorporated despite potentially indicating PSIs.

While the precise molecular mechanisms by which SCFAs influence the gut barrier, brain injury, and bacterial translocation remain to be fully identified and were beyond the scope of this study, three primary pathways have been proposed so far. First, SCFAs act as ligands for G protein-coupled receptors, such as FFAR2, FFAR3, and GPR109A, triggering signalling cascades that modulate immune defences and gut integrity. Second, upon entering cells via monocarboxylate transporters or passive diffusion, SCFAs function as histone deacetylase inhibitors. This epigenetic modulation facilitates an ‘open’ chromatin state, upregulating genes essential for maintaining intestinal integrity and regulating immune responses. Finally, SCFAs serve as a direct metabolic substrate through β-oxidation or conversion into acetyl-CoA, providing the energetic support necessary for cellular activation and tissue repair following injury^24^. In this study, we chose to administer a combination of three naturally occurring SCFAs in proportions mirroring those detected *in vivo* in samples from sudden death patients^76^. Whether the observed protective effects arise from the synergistic action of acetate, butyrate, and propionate, or reflect distinct and complementary contributions of each individual metabolite, has yet to be clarified. Each metabolite exhibits varying affinities for these receptors and unique tissue distribution patterns^8^, suggesting that they may have both overlapping and unique biological effects. Furthermore, whether fluctuations in SCFA levels or receptor expression correlate with the incidence of PSIs or influence their prognosis remains a subject for future research. Finally, as this was a pilot study conducted exclusively in male animals, we acknowledge the limitation of not including female mice. Given the well-documented sexual dimorphism in stroke pathology and immune responses^77^, incorporating both sexes in future research is essential to determine the generalizability of these findings regarding bacterial translocation and to identify potential sex-specific therapeutic outcomes.

In conclusion, the present study advances our understanding of the pathophysiology of PSIs by providing further evidence of systemic and cerebral bacterial translocation following stroke, thereby paving the way for future investigations into the specific bacterial species and mechanisms involved. Our findings highlight the potential of SCFAs as a post-acute, yet effective, novel intervention that not only mitigates brain injury but also preserves intestinal barrier integrity and limits bacterial translocation. By underscoring the pivotal role of the brain-immune-gut axis and microbial metabolites, these results reinforce the promise of SCFA-based therapies as a multimodal strategy to address some of the most lethal and complex complications of stroke.

## Acknowledgements

The authors would like to thank the Cellular Imaging Facility (CIF) staff from the Department of Fundamental Neurosciences (UNIL) for their technical support. We acknowledge support from the Scientific Networks PTI+ NEUROAGING and MICROBIOMA funded by the Consejo Superior de Investigaciones Científicas (CSIC), Spain.

## Sources of funding

This study was funded by the Swiss Science Foundation under FNS 310030_212233 grant to L.H and FNS_10003598 to C.P., and the European Commission ERA-NET NEURON JTC2022 BiotaBB grant and the Biaggi Foundation to L.H. This work was also funded by the Fundación Ramón Areces postdoctoral fellowship and the University of Lausanne under the FBM Transition Grant to J.C-G. The funders played no role in study design, data collection, analysis and interpretation of data, or the writing of this manuscript.

## Disclosures

The authors declare no competing interests.

## Author contributions

J.C-G. conceived and designed the study, conceptualized the experimental framework, conducted the experiments, and performed the formal data analysis. M.M. performed some of the animal procedures and contributed to the brain injury characterization. J-L.R. and J.C-G. analysed the 16S rRNA sequencing data. J.C-G., R.C., S.V., and V.B. performed the flow cytometry experiments and interpreted the resulting data. J.C-G., M.M., J-L.R., R.C., S.V., V.B., L.B., M.P., C.P., L.H., participated in the discussion and interpretation of the findings. J.C-G. drafted the original manuscript. All authors have reviewed, edited, and approved the final version of the manuscript.

## Data availability

All sequencing data were deposited in the NCBI Sequence Read Archive database (PRJNA1509144). Link to reviewers: https://dataview.ncbi.nlm.nih.gov/object/PRJNA1509144?reviewer=2jsu626bs7b2n6b3h6cqneoboi. All other data are available from the corresponding author upon reasonable request.

## Competing interests

All authors declare no financial or non-financial competing interests.

## Notes

### Competing Interest Statement

The authors have declared no competing interest.

### Summary of Updates

In this revised version, we have included new experimental data to provide a deeper understanding of our initial findings. To support this expanded scope, additional studies were conducted, and we have consequently updated the author list to include the new colleagues who contributed to this work.

